# Pattern dynamics on mass-conserved reaction-diffusion compartment model

**DOI:** 10.64898/2026.03.26.714357

**Authors:** Tsubasa Sukekawa, Shin-Ichiro Ei

**Author notes:** **Corresponding author** (S.-I. Ei). E-mail address (T. Sukekawa).

## Abstract

Mass-conserved reaction-diffusion systems are used as mathematical models for various phenomena such as cell polarity. Numerical simulations of this system present transient dynamics in which multiple stripe patterns converge to spatially monotonic patterns. Previous studies indicated that the transient dynamics are driven by a mass conservation law and by variations in the amount of substance contained in each pattern, which we refer to as “pattern flux”. However, it is challenging to mathematically investigate these pattern dynamics. In this study, we introduce a reaction-diffusion compartment model to investigate the pattern dynamics in view of the conservation law and the pattern flux. This model is defined on multiple intervals (compartments), and diffusive couplings are imposed on each boundary of the compartments. Corresponding to the transient dynamics in the original system, we consider the dynamics around stripe patterns in the compartment model. We derive ordinary differential equations describing the pattern dynamics of the compartment model and analyze the existence and stability of equilibria for the reduced ODE with respect to the boundary parameters. For a specific parameter setting, we obtained results consistent with previous studies. Moreover, we present that the stripe patterns in the compartment model are potentially stabilized by changing the parameter, which is not observed in the original system. We expect that the methodology developed in this paper is extendable to various directions, such as membrane-induced pattern control.

## 1 Introduction

In recent years, reaction-diffusion systems with some conservation law have been proposed as mathematical models of various phenomena [6, 12, 10, 15]. Our main concern in this paper is mass-conserved reaction-diffusion systems, which have been used as mathematical models for cell polarity [6, 12, 10]. For these models, it exhibits that stripe patterns appear and converge to spatially monotone patterns such as unimodal patterns (Fig. 1 (a)). In this simulation, the heights of some peaks in the stripe pattern grow, while those of the other peaks decay, and eventually it converges to one peak pattern as in Fig. 1 (a).

**Figure 1.**
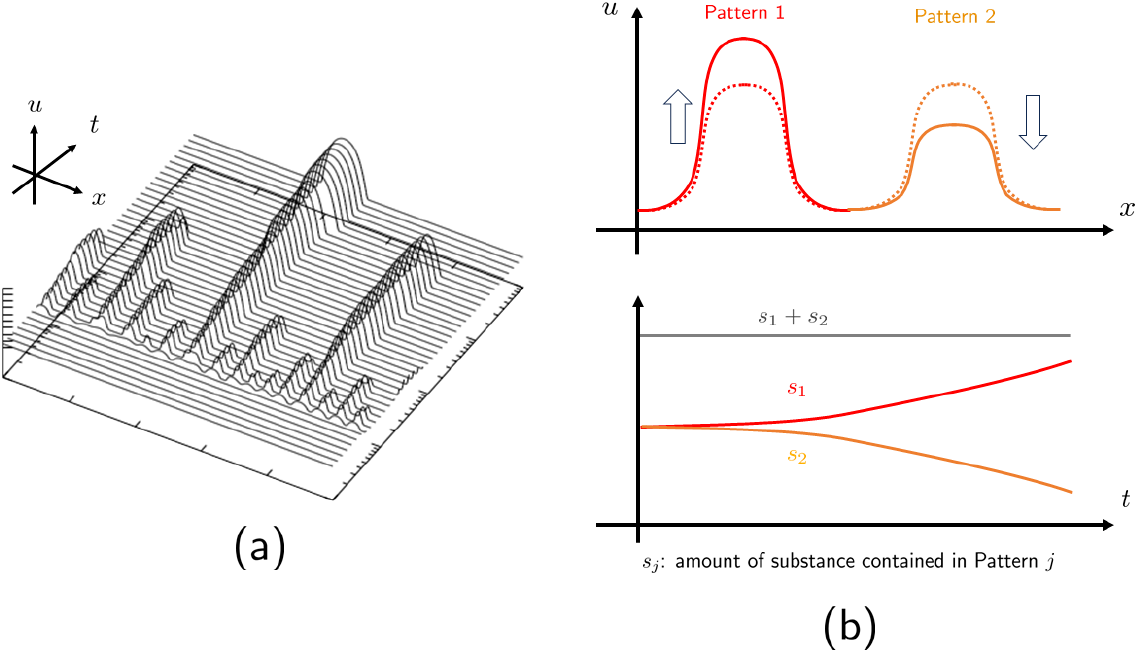
(a) The numerical simulation of the equation (5) with the periodic boundary conditions. *f* (*u, v*) = − *au/*(*u*^2^ + *b*) + *v, a* = 1.0, *b* = 0.1, *d* = 0.02, *τ* = 1.0, *K* = 50.0, *s* = 2.0. The initial condition for the above simulation is the constant stationary solution with a small perturbation. (b) The conceptual image of the pattern flux. In the above image, Pattern 1 grows and Pattern 2 decays. This dynamics can be characterized as the change of the amounts of substance contained in the patterns *s*_*j*_. *s*_1_ and *s*_2_ are increasing and decreasing over time, respectively. However, the total amount of substance *s*_1_ + *s*_2_ is constant due to mass conservation.

Previous studies have been concerned about this growth and decay dynamics [6, 12, 16]. In these studies, the authors were focusing on the amount of substance contained in each pattern constituting the entire stripe pattern(Fig. 1 (a)), and how these amounts change between the patterns. We call this concept “pattern flux” in this study. Namely, if the amount of substance contained in one pattern increases, the amount of substance in the other pattern decreases due to mass conservation (Fig. 1 (b)), then the two-peak pattern becomes unstable.

The previous studies [6, 12, 16] investigated the dynamics with formal mathematical analysis and numerical simulations. The idea of the previous studies is to approximate the stripe pattern using discontinuous functions, and they formally calculate the flux at the points where the function becomes discontinuous.

However, these analyses are not rigorous mathematically. It appears that one of the most difficult parts of the analysis in the previous studies is the discontinuity of the approximating functions. Because the flux in the model equation is defined as the derivative of an unknown function, −*d*_*u*_∂_*x*_*u*, where *u* and *d*_*u*_ represent a concentration of a chemical substance and the diffusion coefficient of the substance, respectively. This means that if we approximate *u* with a discontinuous function, we can not calculate the flux at the points where the function becomes discontinuous.

In this paper, to understand the pattern dynamics of a mass-conserved reaction-diffusion system in view of the mass conservation law and the pattern flux, we introduce a mass-conserved reaction-diffusion compartment model.

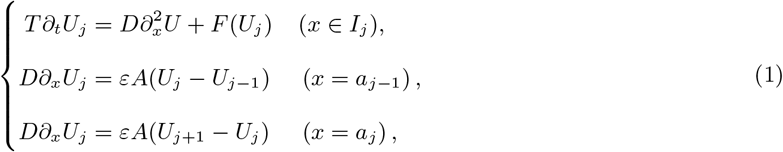

where

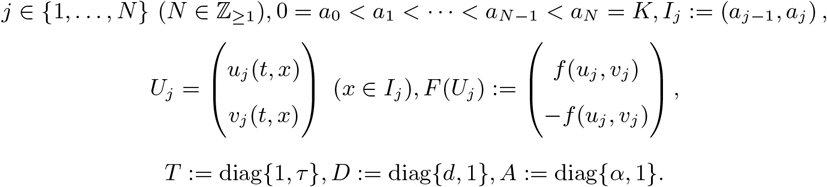

*u*_*j*_, *v*_*j*_ are unknown functions defined on *I*_*j*_. The parameters *α* and *ε* are positive and nonnegative constants, respectively. The equation (1) interpreted as a system of *N* reaction-diffusion systems, where each system is defined on *I*_*j*_. We call each *I*_*j*_ a compartment in this study. The second and third equations in (1) represent boundary conditions, which are diffusive couplings between the compartments. From a physical point of view, the above boundary conditions correspond to a situation where semipermeable membranes partition the interval *I*, and materials are diffusing through the membranes [3]. We note that the parameters *α* and *ε* correspond to the ratio and strength of the diffusive couplings.

In the system (1), *U*_0_(*t*, 0) and *U*_*N* +1_(*t, K*) are determined by boundary conditions at *x* = *a*_0_, *a*_*N*_. When we consider the Neumann boundary conditions at *x* = *a*_0_, *a*_*N*_, we put

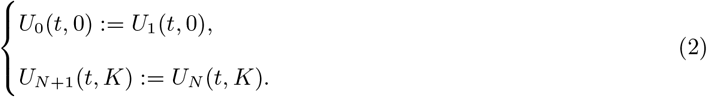

In the case of periodic boundary conditions,

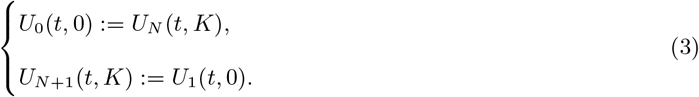

Let us explain the relation between the compartment model (1) and the mass-conserved reaction-diffusion system. A mass-conserved reaction-diffusion system is typically represented as follows:

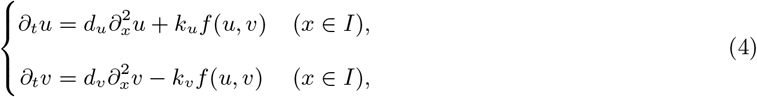

where *I* := (0, *K*). *d*_*u*_, *d*_*v*_, *k*_*u*_, *k*_*v*_ are positive constants. Scaling the variables as 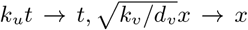, we obtain

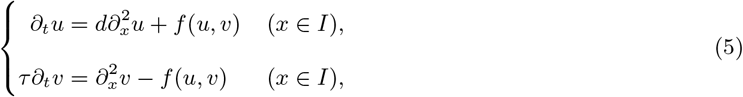

where *d* := (*d*_*u*_*k*_*v*_)*/*(*d*_*v*_ *k*_*u*_), *τ* := *k*_*u*_*/k*_*v*_, and we replaced 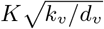 with *K*.

We consider Neumann or periodic boundary conditions for (5).

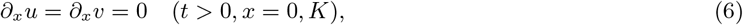

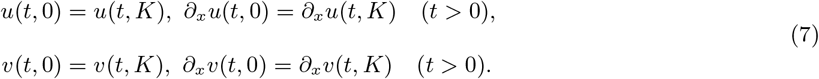

When either the boundary condition (6) or (7) is imposed on the system (5), the following conservation law holds for smooth solutions of (5):

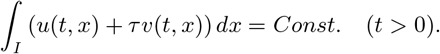

We can check this property by differentiating *∫*_*I*_ (*u* + *τv*)*dx* with the boundary conditions.

To clarify our motivation for this study, we briefly explain the relation between the pattern dynamics observed in Figure 1 and the pattern flux, referring to the previous studies [6, 12, 16]. The authors of the previous studies derive the conditions under which the stripe pattern becomes unstable under the situation where each pattern is close to a stationary solution at the initial time(Figure 1 (b)). Here we consider the two-peak pattern *P* ^2^.

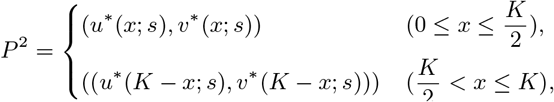

where (*u*^∗^, *v*^∗^) is a stable stationary solution of the equation (5) restricted to the left half 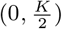 of the original interval (0, *K*) under the periodic boundary conditions, which represents the Pattern 1 in Figure 1 (b), and *s* represent conserved quantity. We note that *P* ^2^ is a stationary solution and the function *du*^∗^(*x*; *s*) + *v*^∗^(*x*; *s*) is constant on 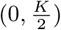. Let us denote *z*(*s*) ≡ *du*^∗^(*x*; *s*) + *v*^∗^(*x*; *s*).

The previous study [16] considered the situation that a solution (*u, v*) to (5) with the periodic boundary conditions, which is approximated at the initial time as follows:

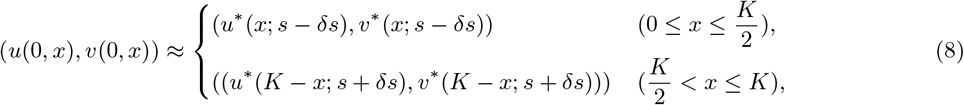

where 0 *< δs* ≪ 1. The initial functions (8) are regarded as the perturbed two-peak solution, namely, the amount of substance on the right half is larger than that on the left half. The previous studies [6, 16] derived the following condition that the two-peak solution becomes unstable:

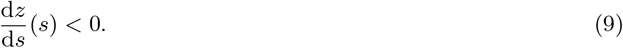

This condition means the total flux of substance at 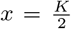 becomes positive [16], namely, the pattern flux is positive, implying the solution tends to leave from the two-peak solution, where both of the left and right intervals have the same amount of substance. In this situation, the pattern dynamics resemble a reverse diffusion, since the substance is transported from the smaller peak to the larger peak.

Our first motivation is to derive the stability criterion (9) in a mathematically rigorous manner. Several previous studies have addressed the stability analysis of equation (5). In some specific conditions, (5) has variational structures, thus the stability of non-constant stationary solutions is established [11, 7, 9]. Recently, singular perturbation techniques have been applied to (5) with reaction terms satisfying bi-stable properties, and the stability of single-layer solutions has been analyzed [8, 5]. However, there seems to be no mathematical analysis from the perspective of the pattern flux, and the formula (9) is not derived mathematically.

The compartment model (1) can be regarded as a modification of the system (5), where the interval *I* is divided into separated *N* intervals *I*_*j*_ (*j* = 1, …, *N*), and a reaction-diffusion system is considered in each interval *I*_*j*_. Same as (5), a following conservation law holds for (1) with (2) or (3):

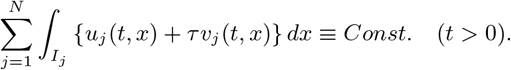

We use the following symbols:

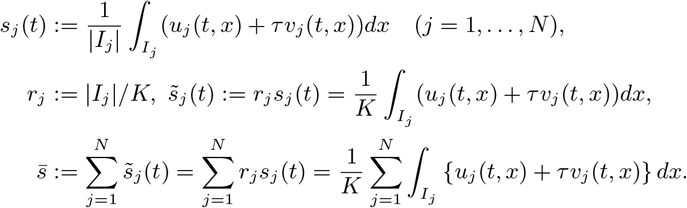

Let us remark that 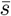 becomes a conserved quantity for (1), but each 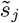 is not conserved generally.

### Remark 1.1.

*When τ* = 1, *the above conservation law can be regarded as the conservation of mass. Then, the function s*_*j*_ *corresponds to the spatial average of the total amount of substances within the compartment I*_*j*_, *whereas* 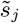 *corresponds to the ratio of the total amount of substances within the compartment I*_*j*_ *and the total length of the system*.

### Remark 1.2.

*We introduce two symbols of functions* 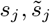, *because each symbol has a different advantage in mathematical analysis. To parametrize a stationary solution of* (1), *s*_*j*_ *is more useful than* 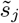 *since we consider the situation ε* = 0 *later and s*_*j*_ *depends only on the equation on I*_*j*_. *When we consider the pattern dynamics of the compartment model with respect to mass conservation, treating* 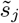 *is simpler, because the sum of* 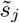 *is conserved even if the lengths of the compartments are not equal; by contrast, the sum of s*_*j*_ *is not conserved in general*.

Here, we summarize the motivations of this study:

### Motivation

1. Understanding the pattern dynamics in Figure 1 and derivation of the stability criteria with respect to the pattern flux.
2. Investigating the pattern dynamics of (1) with respect to *α* and |*I*_*j*_ |

The first motivation has been discussed so far in line with previous studies [6, 12, 16]. The concept of the pattern flux is helpful to understand the pattern dynamics of (5), yet it is challenging to validate mathematically, since the flux is defined as a derivative. In the compartment model (1), the pattern flux is not defined as a derivative but as a difference of functions, see the boundary conditions of (1). Hence, we can treat the pattern flux in a mathematically rigorous manner. In this context, we regard the compartment model (1) as a toy model to (5). This study reproduces the formula (9) through mathematical analysis of the compartment model (1) without formal calculations.

The second motivation concerns the specific pattern dynamics of the compartment model. Initially, we investigated the compartment model in line with the first motivation. However, the compartment model itself can be regarded as a model of a phenomenon including diffusion through a membrane. For example, there is a diffusion phenomenon between cells through a channel connecting membranes, which is called a gap junction, and this diffusion is modeled by boundary conditions of diffusion equations similar to those of the compartment model [2]. From this perspective, each *I*_*j*_ can be regarded as a cell, and the reaction-diffusion systems model the dynamics of chemical concentrations within cells. The parameter *α* and |*I*_*j*_ | correspond to the property of the membrane and the length of each cell, respectively. Mathematical analysis of (1) may lead to understanding such a phenomenon.

Moreover, the analysis of the compartment model may lead to “membrane-induced pattern control”. Since *α*, |*I*_*j*_ | are not present in the original model (5) and are defined in (1) due to the presence of multiple intervals *I*_*j*_, the pattern dynamics in (1) is possibly different from the one in (5) by tuning the new parameters. This means quantitative properties of the pattern dynamics in the original system can be changed by constructing the membranes in the original system. Actually, we numerically observe that the unstable pattern in (5) becomes stable in (1) by choosing the parameter properly.

To accomplish these motivations, we use a regular perturbation technique. When *ε* = 0, the system (5) turns to the non-perturbed system

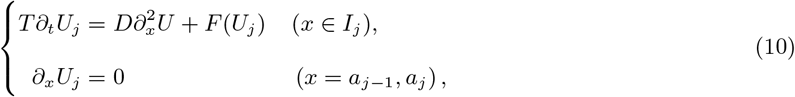

where there is no diffusive coupling between the compartments. The equations (10) represent mass-conserved reaction-diffusion systems defined on *I*_*j*_. Hence, the dynamics of the solutions for (10) is essentially the same as (5) with the Neumann boundary conditions.

Let us denote ***s*** = (*s*_1_, …, *s*_*N*_) ∈ ℝ^*N*^. We denote *P* as a stationary solution of the equation (10) as follows.

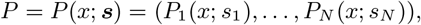

where 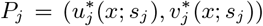 is define on *I*_*j*_ (Fig. 2). Here, *s*_*j*_ is a parameter of *P*_*j*_, which is expressed as 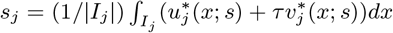. We assume that there exist a vector 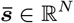 and a constant *δ >* 0 such that a steady-state solution exists for each 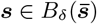, where we denote *B*_*r*_(***p***) := {***s*** ∈ ℝ^*N*^ | ∥***s*** − ***p***∥ *< r*} for any *r >* 0, ***p*** ∈ ℝ^*N*^. Also, we assume that 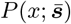 is a stable stationary solution to the equation (10).

**Figure 2.**
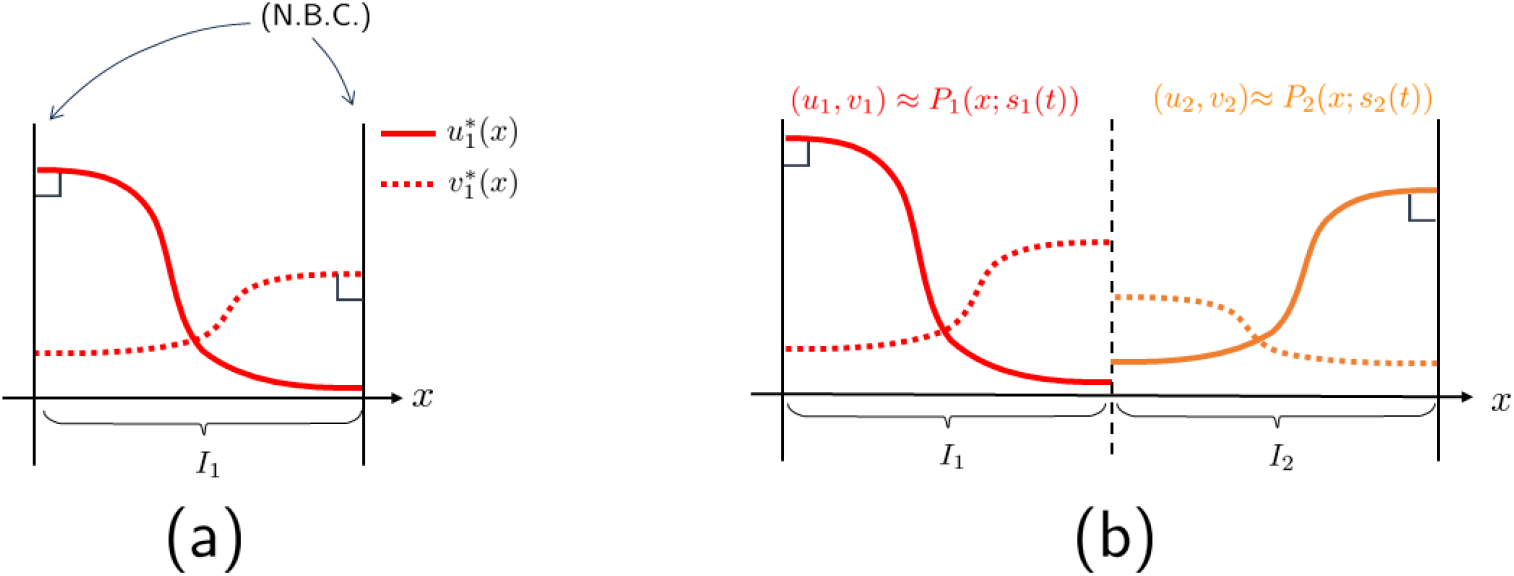
(a) The image of *P*_1_, which is a stationary solution on *I*_1_ with Neumann boundary conditions. (b) The approximation of the solution in the compartment model.

When 0 *< ε* ≪ 1, the boundary conditions in (1) are close to the Neumann boundary conditions. Then we may regard the boundary conditions of (1) as perturbations to *P*. Here we assume that *P* is a stable stationary solution in the non-perturbed system (10). Therefore, if *ε* is sufficiently small and an initial condition (*U*_1_(0, *x*), …, *U*_*N*_ (0, *x*)) is close to *P*, we can expect that the solution of the Cauchy problem for (1) also stay close to the stationary solutions *P* (*x*; ***s***), namely the dynamics of the solution for (1) is essentialy represented as follows (Fig. 2):

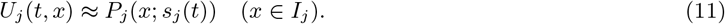

This approximation produces a drastic reduction of the pattern dynamics of (1), because only *s*_*j*_ depends on the time variable *t* in (11). If we understand the time evolution of *s*_*j*_ (*t*), we can appreciate the dominant dynamics, which is represented by (11).

In this paper, we derive the ordinary differential equation of *s*_*j*_ as follows:

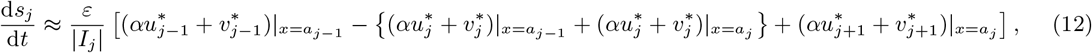

where 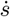 represents time derivative, 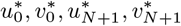 are define by the choice of the boundary conditions (2), (3). We note that the equations (12) can be transformed into the following form:

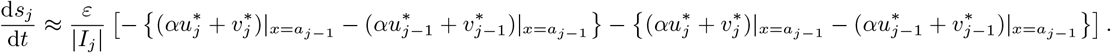

The first and second terms in the square brackets represent the flux of total substance at *x* = *a*_*j*−1_ and *x* = *a*_*j*_, respectively. Namely, the right-hand side of the above equation is the pattern flux of the compartment model.

The equation (12) is a quite general form with respect to the stationary solutions (*u*_*j*_, *v*_*j*_). However, when we obtain an exact form or an approximation of the stationary solutions, we can compute (12) concretely and predict the pattern dynamics of the compartment model. For example, we can compute the following equations with some specific condition on the reaction term and the parameters:

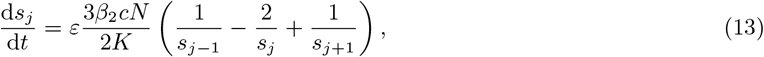

where *β*_2_, *c* are positive constants, see the appendix for the details of the above derivation. We analyze (13) and show the related numerical result in Section 4 and Section 5.

#### Remark 1.3.

*The equation* (13) *has a mathematical structure similar to Ostwald ripening, a coarsening process observed in phase separation phenomena (see [1] and references therein). In this process, it shows that large droplets tend to grow and smaller droplets tend to shrink, showing the similarity to the pattern dynamics of the mass-conserved reaction-diffusion system (Figure 1)*.

In this study, we analyze the existence and stability of equilibrium points in ODE (12). Specifically, the analysis is conducted separately for cases where the lengths of each compartment are equal and where they are unequal. When the compartment lengths are equal 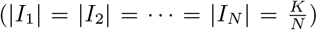, patterns equivalent to the *N* -mode solutions can be handled, corresponding to the stripe patterns in the original model (5). The *N* -mode solution is a stationary solution obtained from a single pattern by reflection extension. This paper considers the *N* -mode solutions in the compartment model and analyzes the pattern dynamics in its vicinity through the ODE (12).

When *α* = *d*, we obtain results equivalent to those concerning the stability of the steady state in the previous study [6]. This suggests that the compartment model qualitatively reproduces the pattern dynamics observed in the original model. Furthermore, the ODE analysis predicted that the *N* -mode solutions could be stable when *α* is sufficiently large under certain conditions. This prediction was verified through numerical simulations. This result indicates that pattern dynamics not observed in the original model are obtained by introducing the membrane parameter.

This paper is organized as follows. In Section 2, we show mathematical preliminaries for the derivation of the ODE of *s*_*j*_. In Section 3, we derive the ODE using the regular perturbation method. In Section 4, we investigate the existence of an equilibrium for the ODE. Also, we analyze the stability of the equilibrium for the ODE. In Section 5, we present numerical simulations. Finally, we summarize the results and discuss the future direction of this paper in Section 6.

## 2 Preliminaries

In this section, we will show the mathematical assumptions in this paper.

Firstly, we consider a stationary solution for the equation (10), which is the unperturbed system of (1). The stationary problem for (10) is here.

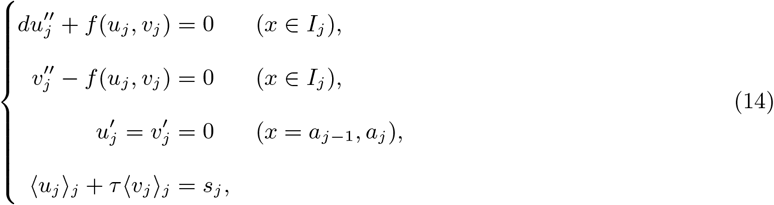

where 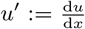 and 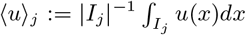. Note that the final equation in (14) corresponds to the conserved quantity of (*u*_*j*_, *v*_*j*_) in (10).

### Assumption 1.

*There exist a vector* 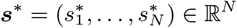 *such that* (14) *has solutions* 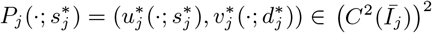, *where* 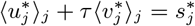.

We sometimes use the term *P*_*j*_ (*s*_*j*_) instead of *P*_*j*_ (·; *s*_*j*_) for convenience.

Let us denote *X*_*j*_ := (*L*^2^(*I*_*j*_))^2^. We define the inner product on the space *X*_*j*_ as follows.

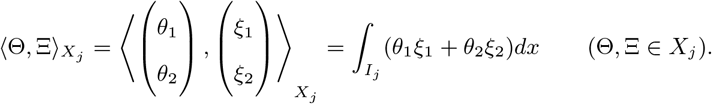

Note that the space *X*_*j*_ become Hilbert space with a norm induced by 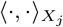. Let us define the norm of *X*_*j*_ as 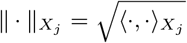. We also define *X* := *X*_1_ × · · · × *X*_*N*_ and 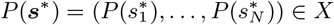.

Here we consider the linearized stability of *P* (***s***^∗^) in (10). For this purpose, we consider linearized stability of 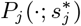, where we here regard *P*_*j*_ as a stationary solution of the equation (5) defined on *I*_*j*_ with the Neumann boundary conditions. Let us define an operator ℒ_*j*_ as linearized operator at 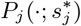.

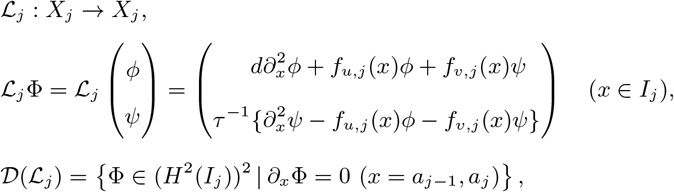

where 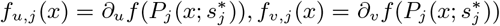.

### Assumption 2.

*There exists η >* 0 *such that σ*(ℒ_*j*_) = {0} ∪ Σ_0_, Σ_*j*_ ⊂ {*λ* ∈ ℂ; *Re λ <* −*η*} *for any j* = 1, …, *N. Moreover*, 0 *is simple eigenvalue of* ℒ_*j*_.

Let 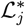 be the adjoint operator of ℒ_*j*_.

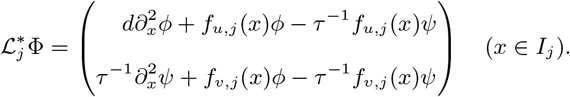

Since ℒ_*j*_ has the simple eigenvalue 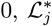 also has it, and let us denote ***a***_*j*_ as a corresponding eigenfunction.

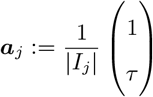

From the Assumption 1,2, the following lemma holds.

### Lemma 2.1.

*Suppose Assumption 1,2 hold. Then there exists a positive constant δ such that there exist maps* 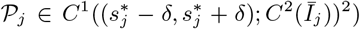, *where* 𝒫_*j*_(*s*_*j*_) *is the solution of* (14) *and* 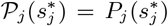. *Let us denote* 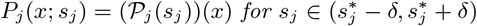.

*Moreover*, 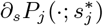 *is a corresponding eigenfunction for* 0 *of* ℒ_*j*_.

*Proof*. see Appendix.

Hereafter, we always consider *P*_*j*_ as same function in the above lemma. Thanks to the above lemma, we can define the functions 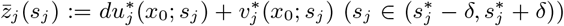, where *x*_0_ is a point of *I*_*j*_ and 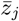 is independent of the choice of *x*_0_. Moreover, 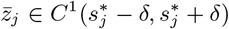. Note that we can prove the existence of the solution curve 𝒫_*j*_ by applying implicit function theorem to the stationary problem (14). We can confirm that 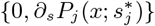 becomes eigen pair of ℒ_*j*_ by differentiating the right-hand side of (14) at 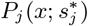 with respect to *s*_*j*_. We refer to Appendix for the detail of the proof.

We remark that *P* (***s***^∗^) is stable stationary solution of (10) under the assumptions 1, 2 (see Appendix). Note that for 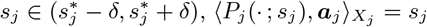. Then, 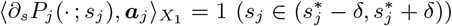.

## 3 Derivation of the equations of *s*_*j*_

In this section, we derive the equation (11). Our strategy to the derivation is applying regular perturbation with regarding *ε* as the strength of the perturbation to the unperturbed system (10). However, we cannot conduct the regular perturbation technique to (1) directly, because the function space, to where the solutions of (1) belong changes with the change of *ε*. This is because the boundary conditions of (1) depend on *ε*.

To avoid this obstacle, we transform the original system (1) to another system with no dependency to *ε* on it’s boundaries. Hereafter, we treat the following system.

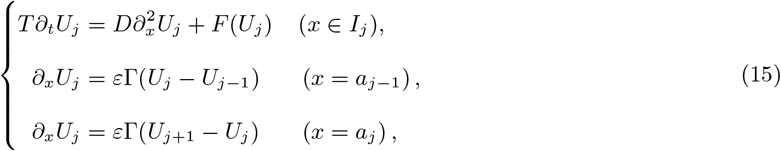

where Γ := diag{*γ*, 1}, *γ* = *α/d*. Obviously, (15) is equivalent formulation to (1).

We consider the following transformation.

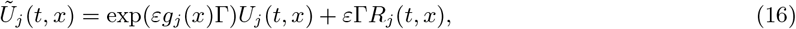

where we suppose *g*_*j*_ ∈ *C*^2^ (*Ī*_*j*_), *R*_*j*_ ∈ (*C*^2^ (*Ī*))^2^. Moreover, we suppose that these functions enjoy the following conditions.

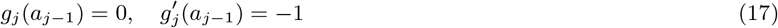

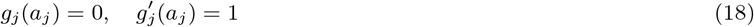

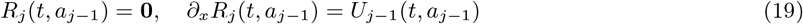

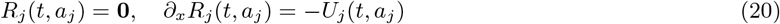

Note that we can construct the functions *g*_*j*_, *R*_*j*_, for instance, by using polynomial functions. Especially, due to the conditions (17) and (18),

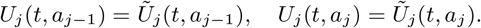

Then, *R*_*j*_ is determined by the boundary values of *Ũ*_*j*_. Moreover, *R*_*j*_ can be regarded as a bilinear map with respect to *Ũ*_*j*−1_, *Ũ*_*j*+1_ when we choose *R*_*j*_ as follows:

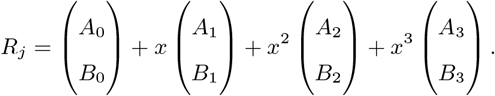

where *A*_*j*_, *B*_*j*_ are unknown constants determined by the boundary conditions (19), (20). Note that 4 boundary conditions are imposed on each component of *R*_*j*_, therefore the constants are uniquely determined by *Ũ*_*j*−1_, *Ũ*_*j*+1_. Also, *A*_*j*_, *B*_*j*_ are linearly dependent on *Ũ*_*j*−1_, *Ũ*_*j*+1_. From these reason, we represent *R*_*j*_ = *R*_*j*_ [*Ũ*_*j*−1_, *Ũ*_*j*+1_].

### Remark 3.1.

*Let us regard U*_*j*−1_, *U*_*j*+1_ *in the boundary conditions of* (15) *as inhomogeneous terms. From this perspective, the roles of the above conditions for g*_*j*_, *R*_*j*_ *are, roughly speaking, eliminating Robin boundary conditions and the inhomogeneous terms, respectively. Then we can impose Neumann boundary condtions on the following equation of Ũ*_*j*_.

We relabel *Ũ*_*j*_ as *U*_*j*_ hereafter. The equation (15) is transformed into the following equation by (16).

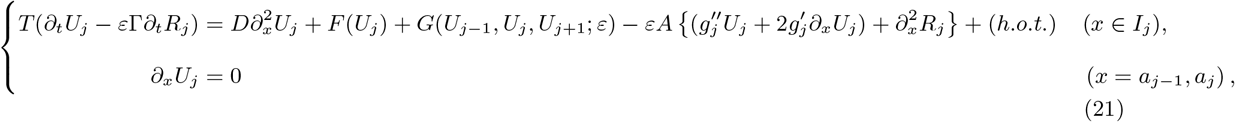

where

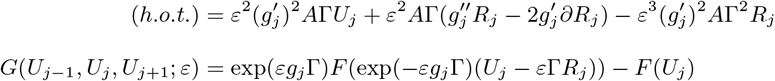

We note that *G*(*U*_*j*−1_, *U*_*j*_, *U*_*j*+1_; 0) = 0.

The boundary conditions for the equation (21) do not depend on *ε*. When *ε* = 0, the equation (21) is same to (10). Hence, we regard (21) as perturbed system to (10) and we conduct regular perturbation method to (21).

Hereafter, we assume 0 *< ε* ≪ 1. Let

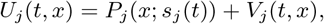

then

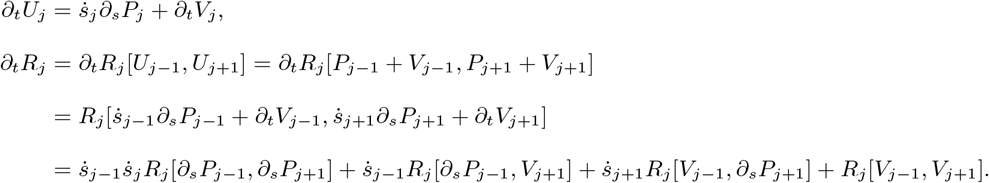

For the relation of ∂_*t*_*R*_*j*_, we used bilinearity of *R*_*j*_. Hence, ∂_*t*_*R*_*j*_ become *O*(*ε*^2^), if all of 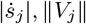 are *O*(*ε*).

When every ∥*V*_*j*_ ∥ = *O*(*ε*),

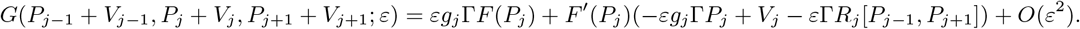

Ignoring the higher order terms and rearranging the equation for *V*_*j*_,

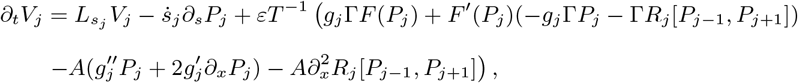

where

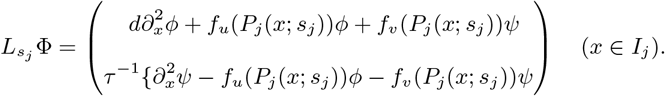

The condition that *V*_*j*_ is bounded in time is

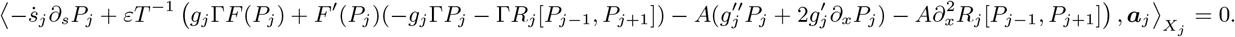

Since 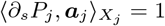,

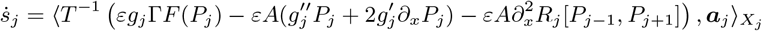

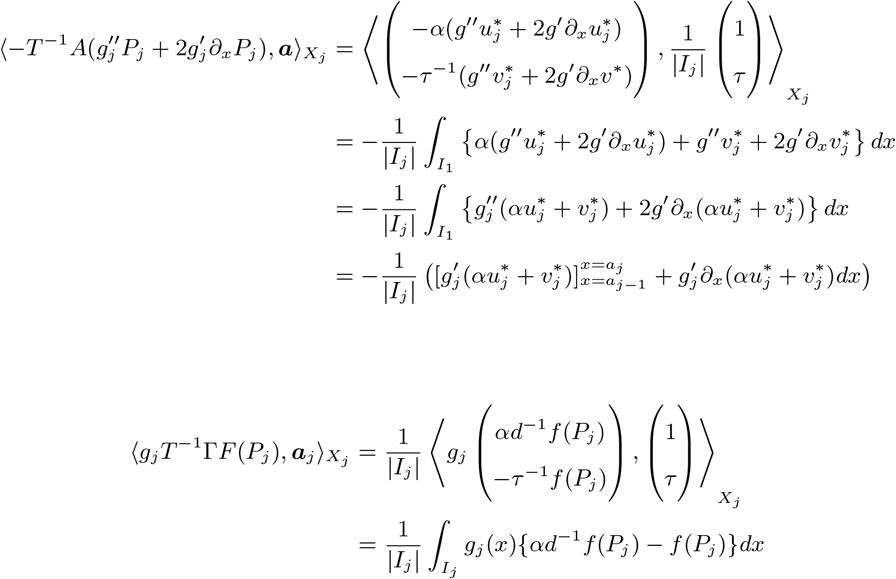

Since 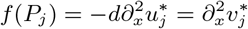,

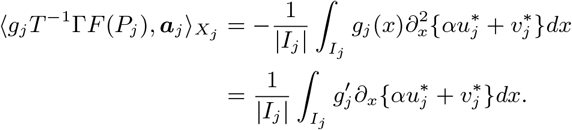

Then,

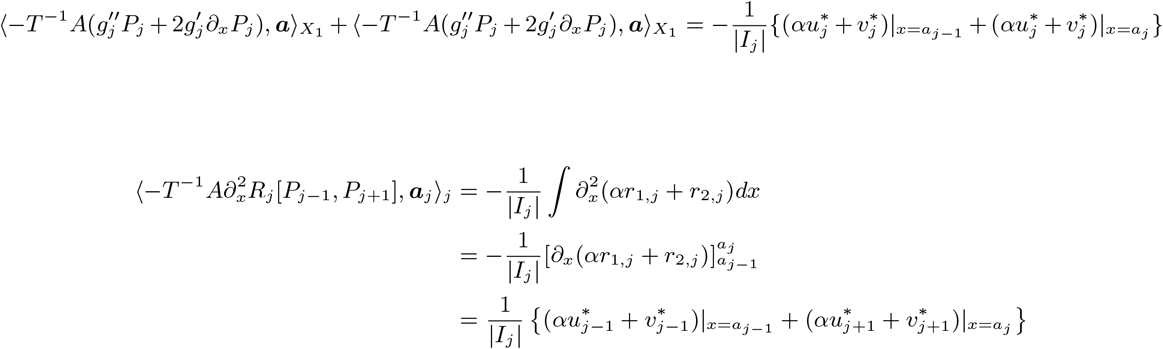

Therefore,

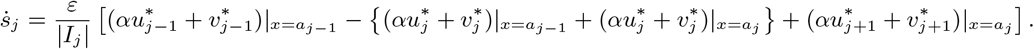

This is the desired equation, same as (11).

## 4 Analysis of ODE

In this section, we analyze the equation (11) in two ways. In the first way, we compare the dynamics of the compartment model with the one of mass-conserved reaction-diffusion system. Especially, to understand “stripe competition” in the mass-conserved system, we consider the dynamics of solution around *N* -mode stationary solution, which is symmetric stationary solution in the mass-conserved system (Fig. 3)and seems to characterize the stripe competition.

**Figure 3.**
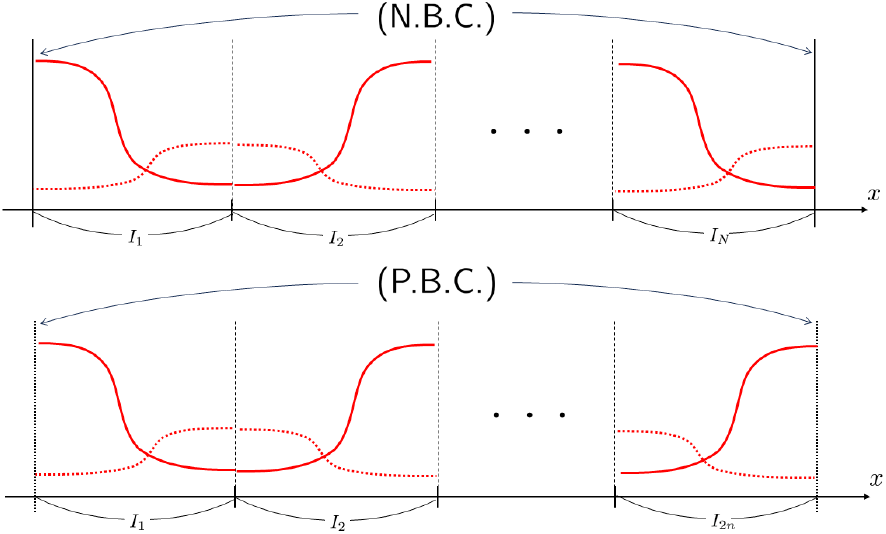
The image of *N* -mode stationary solution

The formula (11) can be represented by more remarkable way. Let us define 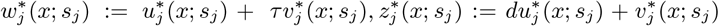. We note that 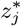 is constant function with respect to the spatial variable *x*. Because *P* is the stationary solution for (10), it follows 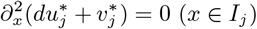. Then, there exists constants *c*_*j*,1_, *c*_*j*,2_ such that 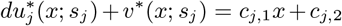. However, *c*_*j*,1_ is 0 due to the Neumann boundary condition. Hereafter, we denote 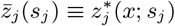.

When 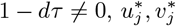 are represented as a linear combination of 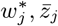 as follows.

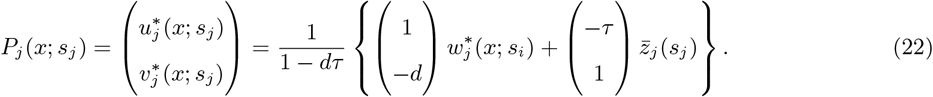

Especially, only 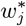 depends on the spatial variable. From these observation, we call 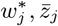 as “wave part” and “constant part” of *P*_*j*_, respectively.

Due to (22),

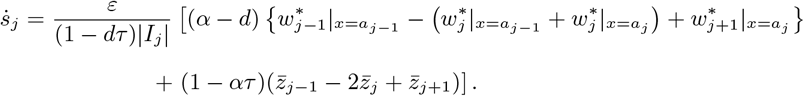

This formula is convenient for investigating the effect of *α* on the dynamics of *s*_*j*_. When *α* = *d*, the terms related to 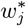, which is the wave part of the stationary solution, disappear. On the contrary, the constant parts 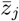 disappear when *α* = *τ*^−1^.

Using the relation 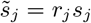, finally we obtain the follwing equation.

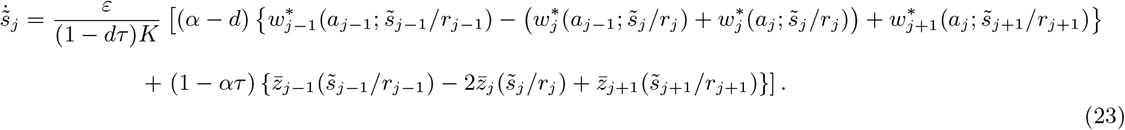

### 4.1 The behavior around *N* -mode stationary solution

Since we consider the *N* -mode stationary solution, we suppose that all compartments have equal length, namely *a*_*j*_ = *jK/N*, |*I*_*j*_ | = *K/N* (*j* = 1, …, *N*). We consider the situation

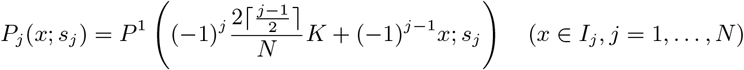

In this case, the equation of *s*_*j*_ is as follows.

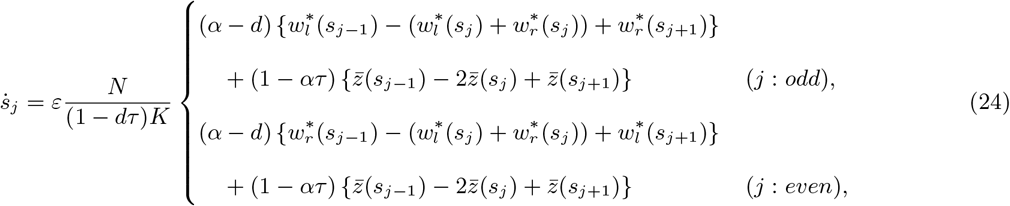

where 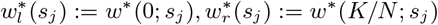.

If we consider the Neumann boundary conditions,

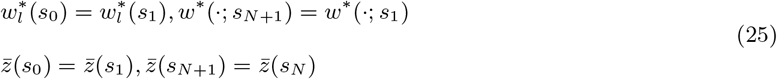

### 4.2 The existence of symmetric equilibrium

It is easy to check that 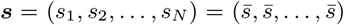 is equilibrium of (24). We note that this equilibrium corresponds to the *N* -mode stationary solution. Let us denote 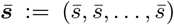. We call 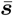 as “symmetric equilibrium” hereafter.

### 4.3 Linearized matrix at symmetric equilibrium

Next, we investigate the dynamics of *s*_*j*_ around the symmetric equilibrium. Let us put 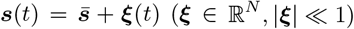. The following is the linearized equation at 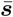 in general representation.

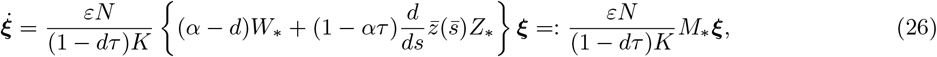

where 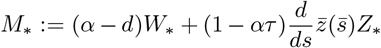. The symbol ∗ distinguish the boundary conditions. Namely, ∗ = N and ∗ = P correspond to the Neumann boundary and the periodic boundary, respectively.

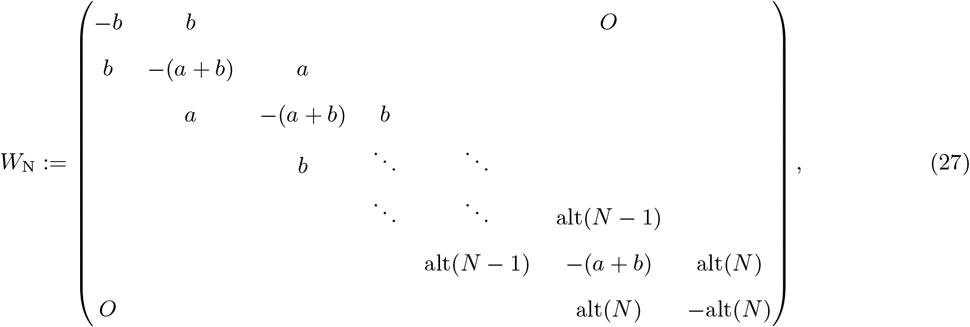

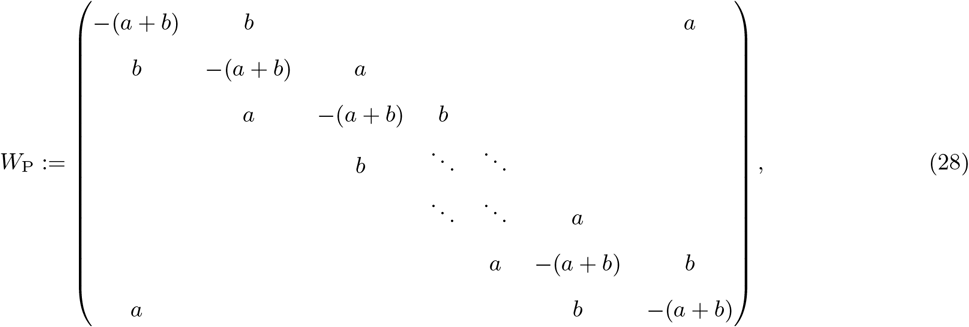

where

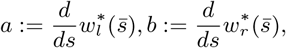

and alt is a alternating function defined as

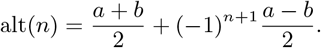

In the case of *Z*_∗_,

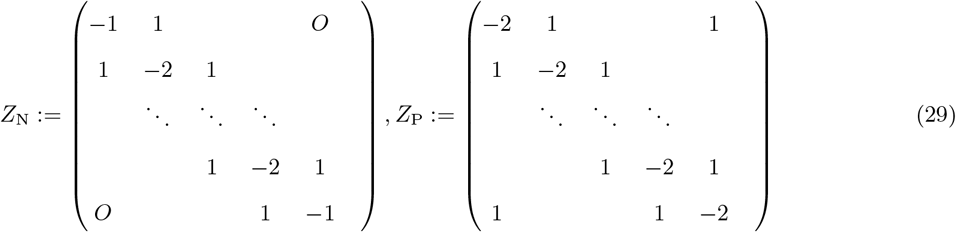

From above, *M*_∗_ is symmetric matrix. Hence, any eigenvalue of *M*_∗_ is real, and the eigenvectors of *M*_∗_ can be chosen to form an orthonormal basis of ℝ^*N*^.

#### 4.3.1 The case: *α* = *d*

When we substitute *α* = *d* to (26), the linearized equation become as follows:

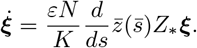

Thus, the stability of the symmetric equilibrium is determined by 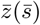 and the eigenvalues of *Z*_∗_. Since *Z*_∗_ is a symmetric matrix, all eigenvalues are real. Then, we denote {*µ*_*m*_} ⊂ ℝ and {*ν*_*m*_} ⊂ ℝ as the sets of eigenvalues of *Z*_N_ and *Z*_P_, respectively. The following lemmas are held:

##### Lemma 4.1.

*Suppose that N* ∈ ℤ_≥1_, *then*

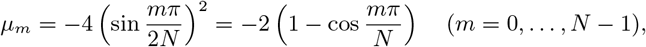

*and an eigenvector* 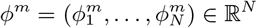 *corresponding to the eigenvalue µ*_*m*_ *is given by the following:*

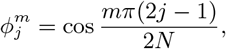

*and also*,

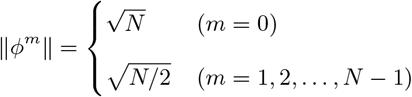

##### Lemma 4.2.

*Suppose that N* = 2*N*^*′*^ (*N*^*′*^ ∈ ℤ_≥1_, *then*

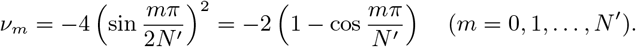

*ν*_0_ *and ν*_*N*_*′ are simple eigenvalues, and the others are semi-simple eigenvalues with multiplicity* 2. *Moreover, let us denote* 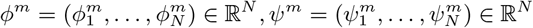 *as follows:*

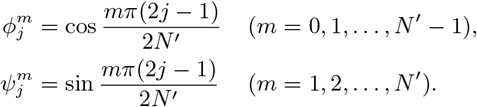

*Then ϕ*^*m*^ *is an eigenvector corresponding to ν*_*m*_ *for m = 0, 1*, …, *N’-1, and ψ*^*m*^ *is an eigenvector corresponding to ν*_*m*_ *for m = 1, 2*, …, *N’*.

*Proof*. See appendix.

Due to the Lemma 4.1 and Lemma 4.2, we obtain the following result:

##### Theorem 4.1.

*Suppose that α* = *d, then the equilibrium* 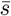 *is unstable if* 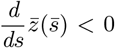. *On the other hand*, 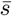 *is stable if* 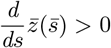.

We note that 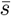 can not be asymptotically stable in general because the zero eigenvalue exists due to the conservation law.

#### 4.3.2 The case: *α ≠ d*

When *α* ≠ *d*, it is difficult to determine the signs of the eigenvalues and to compute the corresponding eigenvectors in general. However, the range of eigenvalues can be estimated using the Gershgorin circle theorem(see, e.g., [4]). Then, we can determine the stability property with respect to the parameter *α*.

##### Theorem 4.2.

*Suppose that* 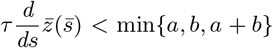. *Then, all eigenvalues of M are negative except for* 0 *for sufficiently large α*.

This result indicates the possibility that the equilibrium 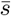 becomes stable when *α* is sufficiently large. In the next section, numerical experiments supporting this result are presented.

Theorem 4.2 may indicate a difference in pattern dynamics between the original mass-conserved reaction-diffusion system (5) and the compartment model (1). To the best of our knowledge, no example is known in which (5) possesses a stable stationary solution with multiple peaks. By contrast, the compartment model (1) may admit such a solution under specific conditions on the parameters related to the boundary conditions. This may suggest that the introduction of new boundaries changes the pattern dynamics of the mass-conserved system. Moreover, the compartment model may also reproduce pattern dynamics similar to those of the original mass-conserved system (see Theorem 4.1 and previous studies [6, 12]). These observations may suggest that the mass-conserved reaction-diffusion compartment model exhibits diverse pattern dynamics.

## 5 Numerical results

In this section, we show the results of the numerical experiments. As a concrete example of the numerical experiments, we consider the following reaction term:

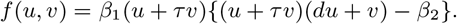

There are exact stationary solutions if we consider the equation (5) on ℝ. The solutions are represented as follows:

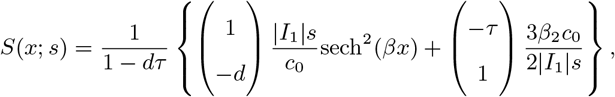

where

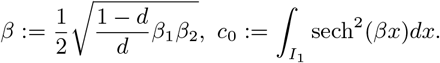

We expect that the above solution becomes a good approximation for the stationary solution on the finite interval *I*_1_ when the length of the interval is large enough. Thus, we use these solutions as the initial condition of the numerical experiments. Hereafter, we use the following parameters:

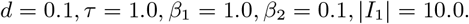

Figure 4 exhibits the numerical simulation for the case of a 2-mode state with the Neumann boundary conditions. The initial conditions and parameters in both simulations are the same, except for the constant *α*. When *α* = *d*, the 2-mode state looks unstable, where the left pattern *u*_1_ decays and the right pattern *u*_2_ grows (See left panels in Figure 4). On the contrary, if we choose *α* = 2.0, the 2-mode state becomes stable (See the right panels in Figure 4).

**Figure 4.**
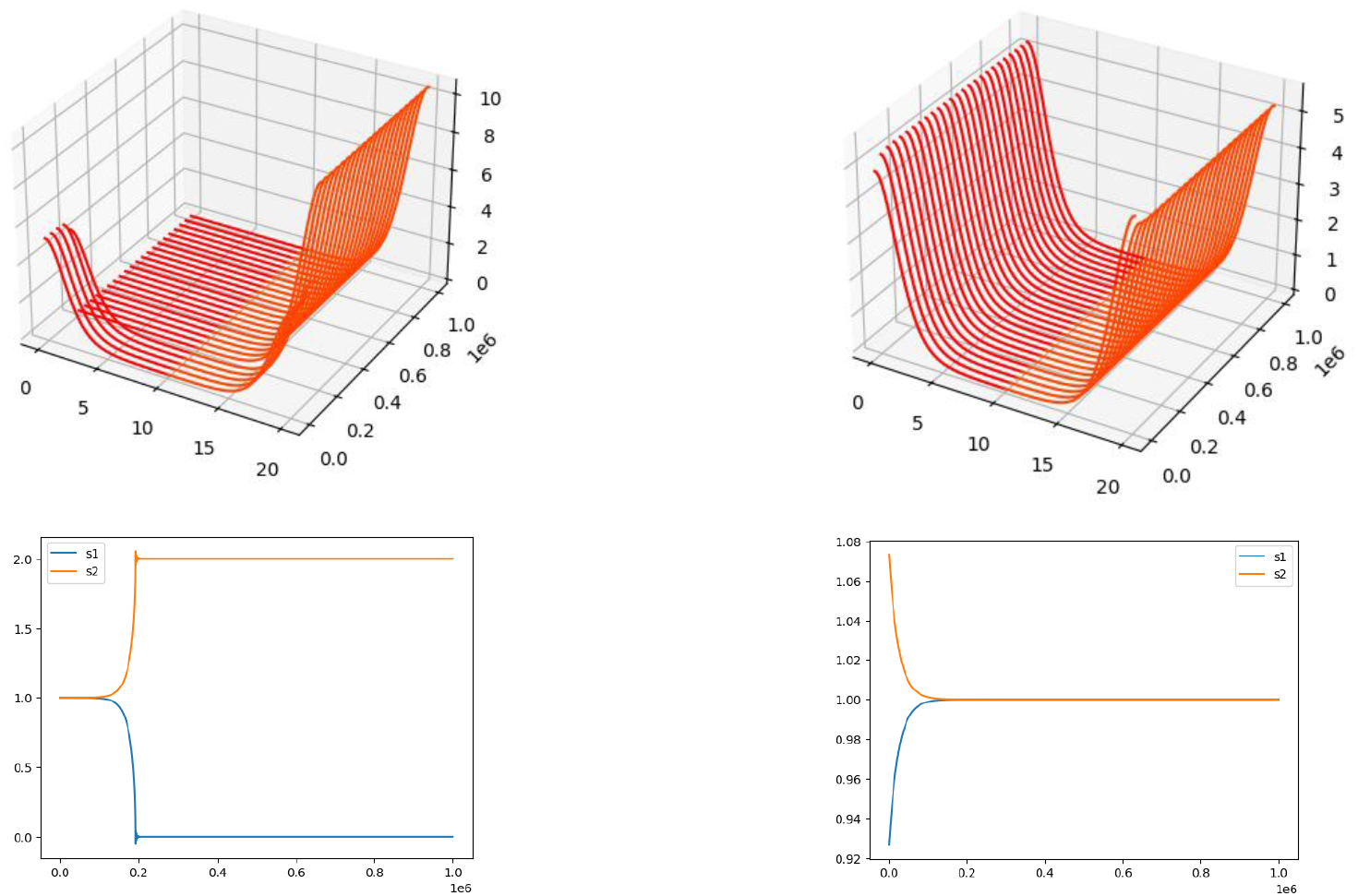
The numerical simulation with the Neumann boundary conditions for the case *N* = 2. The top figures plot *u*_1_(*t, x*), *u*_2_(*t, x*). The bottom figures plot *s*_1_(*t*), *s*_2_(*t*). (Left) *α* = 0.1 = *d*. (Right) *α* = 2.0 *> d*.

This change of the behavior could be explained by Theorem 4.1. Since we have the exact form of the approximated stationary solution, we can represent 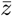 as follows:

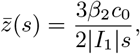

where 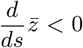. Then Theorem 4.1 implies the 2-mode state is unstable. On the other hand, the constants *a, b* in Theorem 4.2 are represented as follows:

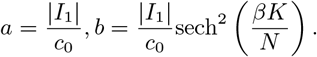

Then, the inequality 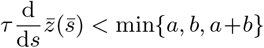 is hold. When we choose *α* large enough, the numerical simulation shows that the 2-mode state is stable (See the right panels in Figure 4).

We note that *S* is originally a stationary solution on ℝ. Thus, it is not certain that these values of 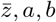 are compatible with the case of the finite interval. However, the results of these numerical experiments appear to be consistent with the consequences of Theorems 4.1 and 4.2.

Figure 5 exhibits the numerical simulation for the case of a 2-mode state with the periodic boundary conditions. The left and right panels show the numerical simulation with *α* = 0.1 and *α* = 2.0, respectively. The stability results are the same as the Neumann boundary case(Figure 4). However, the dynamics’ speed differs significantly from that observed under Neumann boundary conditions. While the growth and decay dynamics cease around *t* = 200, 000 under Neumann boundary conditions, they cease around *t* = 100, 000 under periodic boundary conditions. Intuitively, this is because, under Neumann boundary conditions, interaction occurs at a single point (*x* = *K/*2), whereas under periodic boundary conditions, interaction occurs at two points (*x* = 0 and *x* = *K/*2).

**Figure 5.**
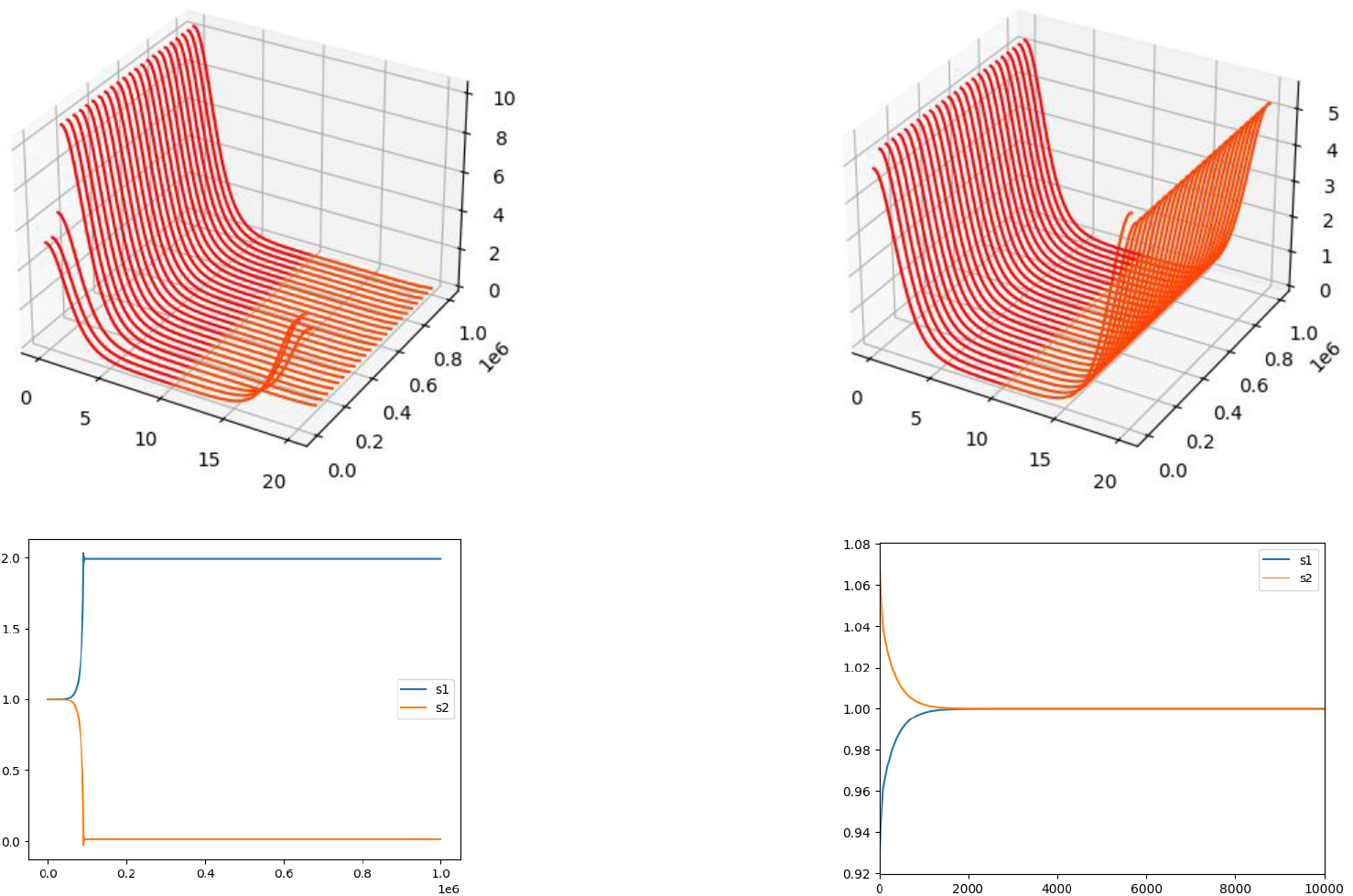
The numerical simulation with the periodic boundary conditions for the case *N* = 2. The top figures plot *u*_1_(*t, x*), *u*_2_(*t, x*). The bottom figures plot *s*_1_(*t*), *s*_2_(*t*). (Left) *α* = 0.1 = *d*. (Right) *α* = 2.0 *> d*.

Furthermore, this difference in speed can be understood in terms of eigenvalues. For *N* = 2,

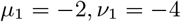

are the eigenvalues with the largest absolute values at *Z*_N_ and *Z*_P_, respectively. When 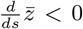, the speed of the dynamics in the corresponding unstable direction is twice that of the Neumann boundary condition. This is consistent with the numerical results presented earlier.

Figure 6 shows the numerical simulation with *N* = 3 under the Neumann boundary conditions. The numerical simulations on the left and right use the same parameters but differ slightly in their initial conditions. Since *α* = *d*, Theorem 4.1 suggests that this 3-mode state is unstable, as in Figure 4. However, its dynamics vary significantly depending on the initial conditions. In the left result, *s*_2_ increases over time, while *s*_1_ and *s*_3_ both decrease. In contrast, the right result shows the exact opposite dynamics, with *s*_1_ and *s*_3_ increasing initially and *s*_2_ decreasing. This difference in dynamics can be understood from the eigenvectors corresponding to the unstable eigenvalues. From Lemma 4.1, when *N* = 3, the eigenvalue with the largest absolute value of *Z*_N_ is *µ*_2_ = −3, and (−1, 2, −1) is a corresponding eigenvector, which represents an unstable direction in the vicinity of the symmetric equilibrium. This vector suggests that, depending on the initial conditions, dynamics may emerge in which *s*_2_ increases and *s*_1_ and *s*_3_ decrease, or *s*_1_ and *s*_3_ increase and *s*_2_ decreases, in the vicinity of the equilibrium point, which is consistent with the numerical results.

**Figure 6.**
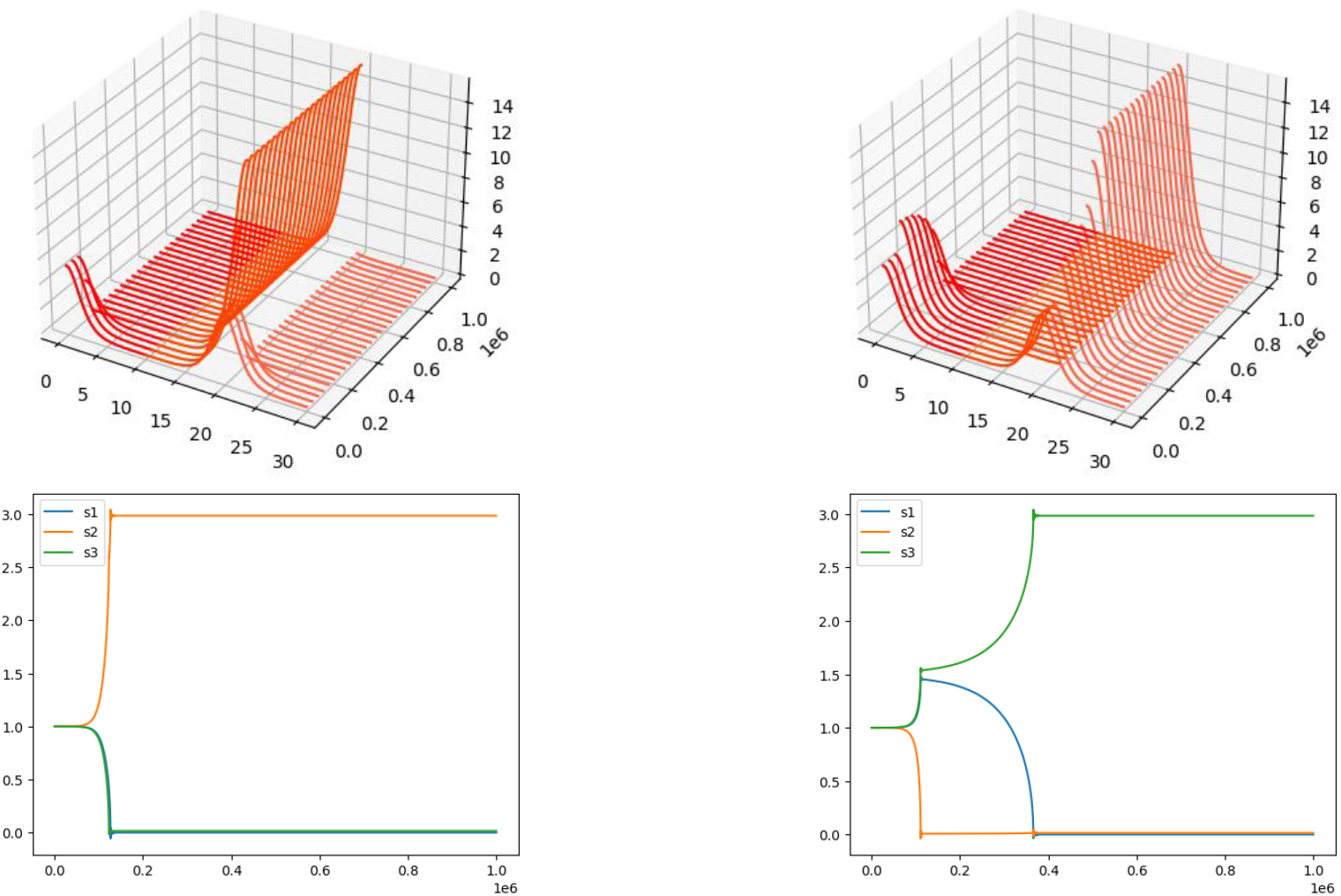
The numerical simulation with the Neumann boundary conditions for the case *N* = 3. The top figures plot *u*_1_(*t, x*), *u*_2_(*t, x*), *u*_3_(*t, x*). The bottom figures plot *s*_1_(*t*), *s*_2_(*t*), *s*_3_(*t*). Here, the same parameters are used on the left and right results: *α* = 0.1, but the initial conditions are slightly changed.

Figure 7 exhibits the numerical simulation for the case of a 4-mode state with the periodic boundary conditions. As same to Figure 6, the numerical simulations on the left and right use the same parameters but have slightly different initial conditions. As in the case of Figure 6, the dynamics in the vicinity of the symmetric equilibrium can be understood from the eigenvectors. From Lemma 4.2, when *N* = 4, the eigenvalue with the largest absolute value of *Z*_P_ is *ν*_2_ = −4, and we can choose (−1, 1, −1, 1) as a corresponding eigenvector. This direction suggests that, depending on the initial conditions, dynamics may emerge in which *s*_2_ and *s*_4_ increases and *s*_1_ and *s*_3_ decrease (Figure 7 (Left)), or *s*_1_ and *s*_3_ increase and *s*_2_ and *s*_4_ decreases (Figure 7 (Right)), in the vicinity of the equilibrium point, which is consistent with the numerical results.

**Figure 7.**
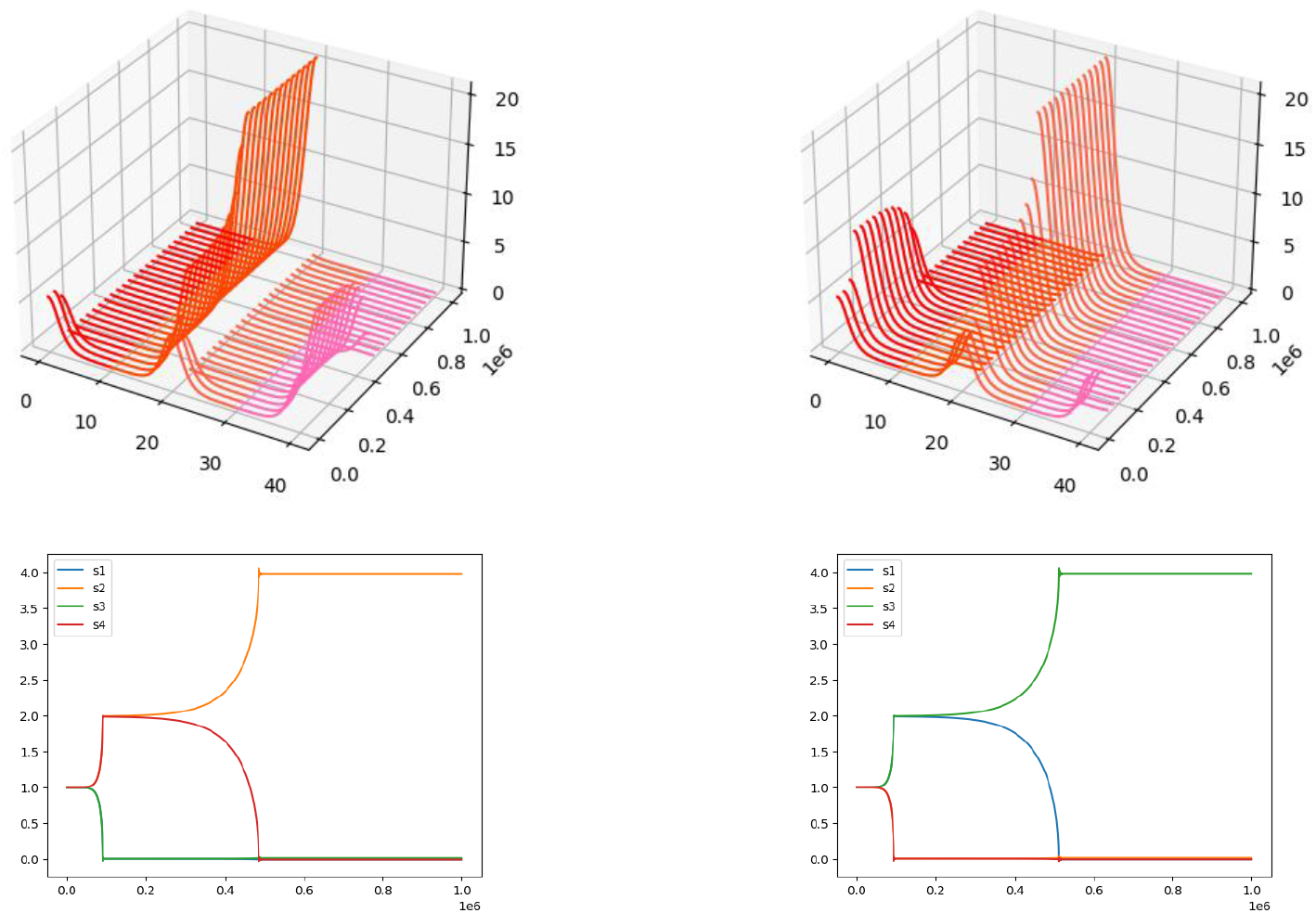
The numerical simulation with the periodic boundary conditions for the case *N* = 4. The top figures plot *u*_1_(*t, x*), *u*_2_(*t, x*), *u*_3_(*t, x*), *u*_4_(*t, x*). The bottom figures plot *s*_1_(*t*), *s*_2_(*t*), *s*_3_(*t*), *s*_4_(*t*). Here, the same parameters are used on the left and right results: *α* = 0.1, but the initial conditions are slightly changed.

## 6 Discussion

In this study, we introduced a mass-conserved reaction-diffusion compartment model to investigate pattern dynamics in a mass-conserved reaction-diffusion system from the perspective of pattern flux. In particular, the transient dynamics shown in Figure 1 (Left) may be interpreted in terms of the mass conservation law and variations in the mass of each pattern (see also Figure 2). To clarify the solution dynamics, particularly near stripe patterns, we formally derived the ODE (12), which describes the time evolution of the integrals of the individual patterns that constitute the stripes, and analyzed the existence and stability of its equilibria. As a result, Theorem 4.1 suggests that when the parameter *α*, representing the diffusion coupling ratio, equals *d*, the instability condition may qualitatively agree with that suggested in previous studies[6].

Furthermore, for the compartment model, it was suggested that patterns unstable in the original mass-conserved reaction-diffusion system may become stable when *α* is taken sufficiently large (Theorem 4.2; Figures,4 and 5). Since the compartment model can be regarded as one that incorporates diffusion through membranes [3, 2], this result may indicate that pattern stability depends on membrane parameters. It is hoped that this result may be applied in future studies to applications such as membrane-based pattern control.

Although this paper has considered only cases with equal compartment lengths, cases with unequal lengths can also be treated theoretically. A possible direction for future research is to investigate how pattern dynamics in compartment models may depend on differences in region size.

We remark that the result of this paper may be varified in the case *ε* is sufficiently small. Another direction of this study is linearized eigenvalue problems to the stationary solutions of the compartment model. From mathematical analysis to these problems, we may understand the dependence of *ε* on the pattern dynamics neighborhood of *N* -mode stationary solutions. Actually, when *N* = 2, this problem is analyzed in the case of Neumann boundary conditions [14]. We can also consider the periodic boundary case. It is analyzed in a forthcoming paper [13].

## Code availability

Numerical codes and all other data are available from the authors on reasonable request.

## Author Contributions

TS and SE launched, designed, and conducted the research. TS excused the research and analyzed the data. SE supervised the research. TS drafted the initial version of the manuscript, and SE is responsible for major revisions. All authors reviewed and revised the final version.

## 7 Appendix

### 7.1 Proof of Lemma 2.1

*Proof*. Let us consider the following map:

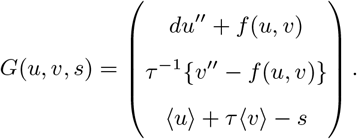

From Assumption 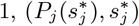 is a zero point of *G*. Let us denote 𝒢_*j*_ as the Fréchet partial derivative of *G* at 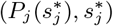 with respect to (*u, v*).

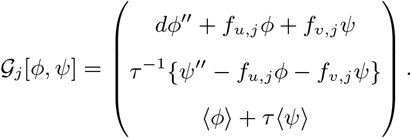

If each 𝒢_*j*_ is bijective for *j* = 1, …, *N*, there exist positive constants *δ*_*j*_ such that there exit curves of stationary solutions 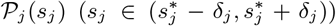 where 𝒫_*j*_ is *C*^1^, due to the implicit function theorem. Let *δ* := min{*δ*_1_, …, *δ*_*N*_ }. Each 𝒫_*j*_ is also *C*^1^ in the interval 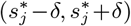, and it is obvious that 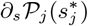 is a eigenfunction of ℒ_*j*_ corresponding to 0. Then the desired theorem holds.

Thus, all we need to prove is that 𝒢_*j*_ is bijective. First, we prove that 𝒢_*j*_ is injective. Suppose that

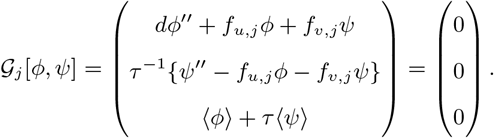

Due to the Assumption 2, (*ϕ, ψ*) is represent as

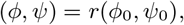

where (*ϕ*_0_, *ψ*_0_) is a nonzero eigenfunction for 0 of ℒ_*j*_. From the simlicity of 0, the adjoint operator 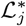 also has simple eigenvalue 0, and ***a*** = |*I*_*j*_ |^−1^(1, *τ*) is a corresponding eigenfunction. Then,

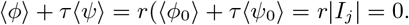

This conclude than (*ϕ, ψ*) = (0, 0). Hence, 𝒢_*j*_ is injective.

Second, we prove surjectivity of 𝒢_*j*_. Fix any (*ϕ*_1_, *ψ*_1_, *s*_1_). We consider the following equations:

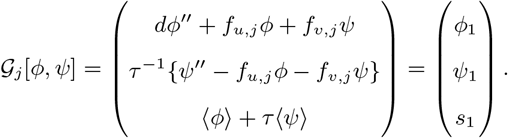

Due to the first and second equations, ⟨*ϕ*_1_⟩ + *τ* ⟨*ψ*_1_⟩ = 0. Thus, there exists a solution (*ϕ, ψ*) to these equations, and it is represented as follows:

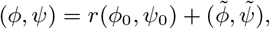

where 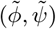 is a peculiar solution to these equations. From the third equation,

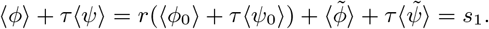

Then if we choose

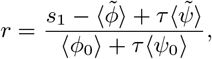

(*ϕ, ψ*) become the solution to the above equations. Hence, 𝒢_*j*_ is surjective.

### 7.2 Proof of Lemma 4.1

*Proof*. We note that 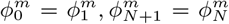 for each *m* ∈ ℤ. Thus, the *j*-th component of the vector *Z*_N_*ϕ*^*m*^ is represented as follows:

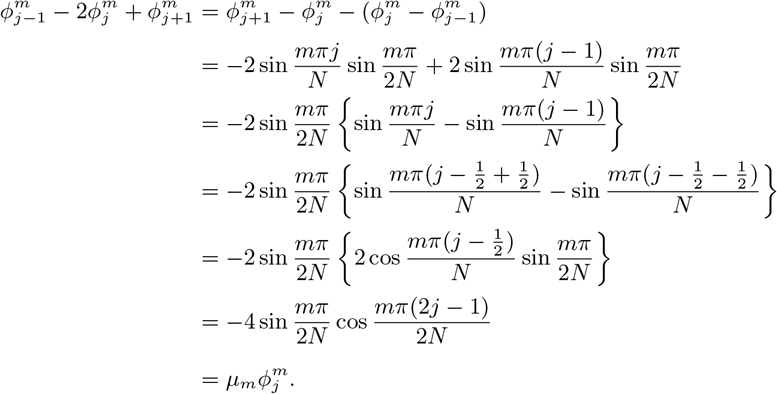

Hence, {*µ*_*m*_, *ϕ*^*m*^} is the eigen pair of *Z*_N_.

Next, we prove the formula of ∥*ϕ*^*m*^∥. It is obvious for the case *m* = 0. When *m* ≠ 0,

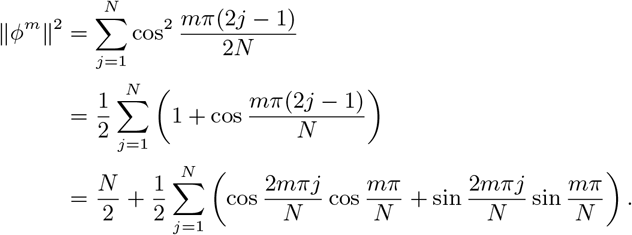

Here, we denote 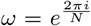, where 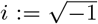. Note the fact that 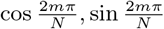 are real and imaginary part of *ω*^*m*^, respectively, and when *m* ∈ {1, 2, …, *N* − 1},

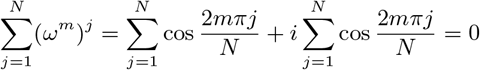

due to the property of *N* -th roots of 1. Then, 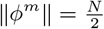 holds for *m* = 1, …, *N* − 1.

### 7.3 Proof of Lemma 4.2

*Proof*. Note that 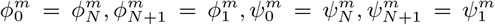. Then, the *i*-th components of the vectors *Z*_P_*ϕ*^*m*^, *Z*_P_*ψ*^*m*^ are represented as follows, respectively:

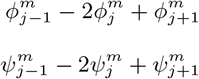

Due to the proof of Lemma 4.1, {*ν*_*m*_, *ϕ*^*m*^} become the eigenpair. Because we can prove that {*ν*_*m*_, *ψ*^*m*^} is also eigenpair by the same line, we skip the detail of the proof.

Next, we prove that the dimension of the eigenspace for *ν*_*m*_ (*m* = 1, …, *N*^*′*^ − 1) is exactly 2. Let us compute the inner product ⟨*ϕ*^*m*^, *ψ*^*m*^⟩ for *m* = 1, …, *N*^*′*^ − 1.

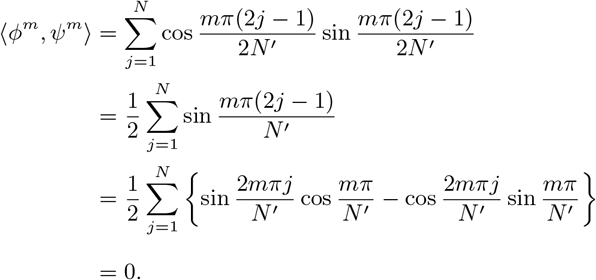

The last equality holds from the facts

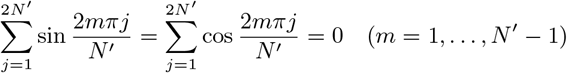

due to the property of *N*^*′*^-th root of 1. Hence, *ϕ*^*m*^, *ψ*^*m*^ are linearly independent, concluding the dimension of the eigenspace for *ν*_*m*_ (*m* = 1, …, *N*^*′*^ − 1) is 2.

